# Inferring the demographic history of coppery titi monkeys (*Plecturocebus cupreus*) from high-quality, whole-genome, population-level data

**DOI:** 10.64898/2026.01.09.698678

**Authors:** John W. Terbot, Vivak Soni, Cyril J. Versoza, Karen L. Bales, Susanne P. Pfeifer, Jeffrey D. Jensen

**Affiliations:** Center for Evolution and Medicine, School of Life Sciences, Arizona State University, Tempe, AZ, USA; Department of Psychology, University of California, Davis, CA, USA; California National Primate Research Center, University of California, Davis, CA, USA

**Keywords:** primate, haplorrhine, Pitheciidae, demographic inference, population history, population genetics

## Abstract

Despite being an important biomedical model species for social behavior, the natural population history of the coppery titi monkey (*Plecturocebus cupreus*) remains largely uncharacterized, in part due to the scarcity of genomic resources available for the species. Apart from the inherent interest in the demographic dynamics of this abundant platyrrhine native to the Amazon forest of Brazil and Peru, this quantification will also serve as a central component of future genotype-to-phenotype studies, given the ability of historical population size change and structure to generate genetic associations. In this study, we deep-sequenced the genomes of six unrelated individuals and inferred a baseline demographic model based on observed levels and patterns of variation in the non-coding regions of the genome. In characterizing these demographic dynamics, we found that estimated population size changes correspond well to previously described speciation times as well as to large-scale climatic changes relating to glaciation patterns.

## Introduction

The coppery titi monkey (*Plecturocebus cupreus*) is a relatively abundant and diminutive platyrrhine species primarily found in Western Brazil and Eastern Peru (Heymann et al. 2021). Within this area, they are mainly restricted to habitats of lowland forests subject to periodic seasonal flooding, their home ranges are generally non-overlapping, and offspring leave their family units around 2 to 3 years of age (Mason 1966; Dolotovskaya et al. 2020; Conley et al. 2022). With a lifespan of over 20 years (Zablocki-Thomas et al. 2023), *P. cupreus* is notable for being characterized by a monogamous, pair-bonded mating system. Given that social monogamy is a relatively rare social system in mammals, and pair-bonding is rarer still, this species has become an important model to study the neurobiology of social behavior (e.g., Bales et al. 2007; Lau et al. 2024; and see Bales et al. 2021). For example, in species characterized by this social system, visually-associated brain regions have been found to contain a high density of receptors for the social hormone oxytocin (Freeman et al. 2014), suggesting a key role of vision in governing this behavior (Baldwin and Krubitzer 2018). This primate is thus also of great interest in a comparative framework for the study of complex human social behavior and attachment, and considerable work has focused on the systems of dopamine, oxytocin, and arginine vasopressin in the hypothalamus, globus pallidum, and other limbic and cortical regions in this regard (Feldman 2017; Fischer et al. 2019).

*P. cupreus* is a member of the Western Amazon clade of the *moloch* group alongside *P. moloch*, *P. brunneus*, *P. dubius*, and *P. caligatus*. The *moloch* group is believed to have diverged around 3.78 million years ago (mya) during the drying of the Pebas system (a large lake and floodplain covering much of what is now the Western Amazon) in the late Neogene period (Byrne et al. 2018). Further changes to the landscape during the early Pleistocene are thought to have led to the additional split in this group between Western and Eastern Amazonian clades around 1.95 to 3.44 mya. This was followed by multiple proposed speciation events including a split between the clade containing *P. cupreus* and the clade containing *P. caligatus* around 1.44 to 1.95 mya, as well as subsequent subspecific divisions in *P. cupreus* (Byrne et al. 2016, 2018; Byrne 2017).

To complement these previous genus-level estimates, and given the biomedical importance of *P. cupreus*, we here present a detailed demographic analysis of the species based on novel, whole-genome, high-quality sequencing data from six unrelated individuals. In order to perform this estimation, we implemented two of the most commonly used inference approaches — *δaδi* (Gutenkunst et al. 2009) and fastsimcoal2 (Excoffier et al. 2013) — both of which rely on fitting models, and parameters underlying those models, to the empirically observed site frequency spectrum (SFS). By utilizing patterns of genetic diversity at putatively neutral, intergenic sites of the genome sufficiently distant from functional regions to avoid the confounding effects of background selection (Soni and Jensen 2025; Soni, Versoza et al. 2025a; Terbot et al. 2025a, Terbot et al. 2025b), the well-fitting demographic model described here provides new insights into the population history of this species. Furthermore, this demographic inference will also serve as a necessary neutral baseline model for future genomic scans of episodic positive or balancing selection (Poh et al. 2014; Johri et al. 2022; Jensen 2023; Soni and Jensen 2024; Soni, Terbot et al. 2025; Soni et al. 2025), related to, for example, the underlying genetic modifications governing their social behavior, as well as to future association studies seeking to connect genotypes to well-studied phenotypes. Our results suggest that the sample of this study was derived from a single sub-population, and we found evidence of a size change history correlated with past speciation events as well as glaciation / interglacial climatic patterns. As these climatic shifts were associated with dryer and cooler weather, it is likely that these fluctuations were related to the abundance of dry or temporarily flooded forest habitats of the sort preferred by the species to this day.

## Materials and Methods

### Ethics statement

This study was performed in compliance with all regulations regarding the care and use of captive primates, including the NIH Guidelines for the Care and Use of Animals and the American Society of Primatologists’ Guidelines for the Ethical Treatment of Nonhuman Primates. Procedures were approved by the UC-Davis Institutional Animal Care and Use Committee (protocol 22523).

### Samples and sequencing

We collected blood samples from six unrelated coppery titi monkeys housed at the California National Primate Research Center (CNPRC) during routine veterinary care. DNA extracted from each sample with the PAXgene Blood DNA System (Qiagen, Hilden, Germany) was used to prepare individual libraries following a PCR-free protocol. Libraries were sequenced on an Illumina NovaSeq 6000 (Illumina, San Diego, CA, USA) with a 2 × 150 bp sequencing configuration with a target of ∼40× coverage.

### Whole genome sequencing alignment and variant calling

We removed adapters from the sequencing reads using TrimGalore v.0.6.10 (https://github.com/FelixKrueger/TrimGalore) with Cutadapt v.4.9 (Martin 2011) built-in and subsequently mapped the reads to the reference genome sequence of the species, PleCup_hybrid (NCBI GenBank ID: GCA_040437455.1; Pfeifer et al. 2024), using BWA-MEM v.0.7.17 (Li 2013).

Following best practices (Pfeifer 2017), we marked duplicate reads using the *MarkDuplicates* function implemented in the Genome Analysis Toolkit (GATK) v.4.5.0.0 (van der Auwera and O’Connor 2020) to remove technical duplicates from library preparation and sequencing. Additionally, to avoid systematic biases in the base quality scores emitted by the sequencer, we recalibrated base scores using GATK’s *BaseRecalibrator* and *ApplyBQSR* functions together with high-quality training data from pedigreed individuals of the species (Versoza et al. 2026a). Afterward, we used GATK’s *HaplotypeCaller* function to first call both variant and invariant sites (’*-ERC* BP_RESOLUTION’) from the high-quality recalibrated reads (’*--minimum-mapping-quality* 40’) of each sample (with the PCR error correction disabled as a PCR-free library protocol was followed during sequencing: ’*-pcr-indel-model* NONE’), merged individual calls using GATK’s *CombineVCF* function, and then jointly genotyped all samples using GATK’s *GenotypeGVCFs* function with the ’*-all-sites*’ flag enabled. We limited the call set to autosomal, biallelic variant (’*--restrict-alleles-to* BIALLELIC *--select-type-to-include* SNP’) and monoallelic invariant (’*--select-type-to-include* NO_VARIATION’) sites genotyped in all individuals (’*AN =*= 12’) using the *SelectVariants* function implemented in GATK v.4.2.6.1 (van der Auwera and O’Connor 2020) and then filtered sites using the *VariantFiltration* function according to the recommendations of the developers (’*-filter* QD<2.0 *--filter-name* QD2 *-filter* QUAL<30.0 *--filter-name* QUAL30 *-filter* SOR>3.0 *--filter-name* SOR3 *-filter* FS>60.0 *--filter-name* FS60 *-filter* MQ<40.0 *--filter-name* MQ40*-filter* MQRankSum<–12.5 *--filter-name* MQRankSum-12.5 *-filter* ReadPosRankSum<–8.0 *--filter-name* ReadPosRankSum-8’). Additionally, we removed any sites overlapping repetitive / low- complexity regions using the *intersect* function implemented in BEDTools v.2.30 (Quinlan and Hall 2010) based on the annotations of the coppery titi monkey reference genome (Pfeifer et al. 2024) as well as those exhibiting extreme coverage (defined here as sites with less than 0.5, or more than 2, times the mean sequencing depth of a sample) using the GATK *SelectVariants* function, as such regions tend to be prone to mis-mapping, variant calling, and genotyping errors.

### Population genomic data for demographic inference

As both direct and background selection can bias the inference of demographic history (Ewing and Jensen 2014, 2016; Johri et al. 2020, 2021; Charlesworth and Jensen 2021, 2024), we followed the recommendations of Johri et al. (2020, 2023) and restricted the high-quality call set to putatively neutral genomic regions. To this end, we used BEDTools *intersect* v.2.30 (Quinlan and Hall 2010) to exclude both sites overlapping with protein-coding sequence (Pfeifer et al. 2024) or non-coding regulatory sequence elements under selective constraint across primates (Kuderna et al. 2024), thus controlling for the effects of purifying selection; we also excluded sites located within 10 kb of exons, thus controlling for the effects of background selection. To improve inference, these putatively neutral sites were phased using BEAGLE v.5.5 (Browning et al. 2021).

### Estimating population structure

To determine the number of sub-populations present among the titi monkeys sequenced in this study (six diploid individuals, or 12 haploids), we analyzed putatively neutral data from each chromosome independently and for the entire genome combined using fastSTRUCTURE v.1.0 (Raj et al. 2014) — a software that uses a variational Bayesian framework to implement the optimization algorithms from the STRUCTURE program (Pritchard et al. 2000; Falush et al. 2003). Specifically, we performed analyses for values of *k* (number of demes) from 1 to 5 and selected the optimal number of demes based on the value of *k* that maximized the marginal likelihood.

### Inferring the population size-change history

Our fastSTRUCTURE results strongly favored a single deme model. Thus, we analyzed single-population demographic models with six different model structures (0 to 5 size change events) using fastsimcoal2 v.2.8.0.0 (Excoffier et al. 2013, 2021; Marchi et al. 2024) — a software that compares the composite likelihood scores of SFS simulated under various parameters to that of the empirical SFS, and then selects the best-fitting parameters by maximizing this composite likelihood while minimizing the difference between the simulated and empirical scores. For all fastsimcoal2 models, we bounded population sizes between 1,000 and 10,000,000 haploid genomes (i.e., 500 to 5,000,000 diploid individuals), and initially set the timing of events to between 1 to 300,000 generations for the single-event model and to between 1 to 500,000 generations for all other models, though the upper limit for event timing was not bounded. For each of the six model structures (0 to 5 size change events), we ran 500 replicates based on the folded SFS, each with 150,000 rounds of simulation per parameter and 50 maximization cycles, re-setting the parameter search after 10 consecutive failed cycles to improve the likelihood score (’*-n*150000 *-L* 50 *-y* 10 -M *-u -m*’). We assumed a neutral mutation rate of 4.88e-9 /site/generation and a recombination rate of 1e-8 events/site/generation. Notably, this mutation rate was based on pedigree-based (i.e., direct) estimation (Versoza et al. 2026a). In order to examine the impact of uncertainty in this underlying mutation rate, we also examined demographic model scaling at a rate of 1.07e-8 /site/generation for comparison, as suggested by divergence-based (i.e., indirect) estimation using a six-year generation time and an estimated divergence time of 1.16 million years between the *cupreus* clade and the *caligatus* clade within the Western *moloch* group (Soni, Versoza, et al. 2026). This range of rates is comparable to other estimates obtained from non-human primates (see the reviews of Tran and Pfeifer 2018; Chintalapati and Moorjani 2020). Similarly, the recombination rate was based on a recent estimate obtained from pedigree data in the species (Versoza et al. 2026b), which too resembles rates observed in other non-human primates (e.g., Versoza, Weiss et al. 2024; Soni, Versoza et al. 2025b; Versoza, Lloret-Villas et al. 2025).

Following the analyses using fastsimcoal2, we sought to confirm the estimated demographic history using *δaδi* v.2.1.0 (Gutenkunst et al. 2009) — a demographic inference software that optimizes parameters using a diffusion approximation approach and a continuous approximation of the expected SFS under particular demographic models for comparison to the observed SFS. We considered seven single-population models: the standard neutral model, as well as 2-epoch (a single instantaneous population size change), 3-epoch (two instantaneous population size changes), 4-epoch (three instantaneous population size changes), 5-epoch (four instantaneous size changes), growth (exponential population size change), and bottlegrowth (an instantaneous population size change followed by an exponential population size change) models. For each model, we performed inference 100 times using the *dadi.Inference.optimize* function, with a maximum of 300 iterations (*maxiter*) and [30, 50, 70] grid points (*pts*) (i.e., larger than the haploid sample size of *n* = 12), together with the *δaδi*’s *perturb_params* function to adjust the starting parameters two-fold up or down, within the parameter bounds defined in Supplementary Table 1. As *δaδi* infers the ancestral population size from *θ*, we calculated all other parameters relative to this population size.

### Distinguishing between estimated demographic histories

We determined the best-fitting fastsimcoal2 model by first comparing the composite likelihood scores of the best parameterized version of each model and then assessing the number of replicates for each model that represented improvements over the best-performing, next simplest model. Additionally, we performed a final screening by quantifying the consistency of the specific parameters estimated for each model’s best performing replicates using z-scores, calculated based on the mean and standard deviation of each parameter across those replicates. For the *δaδi* models, we selected the model with the highest log-likelihood score across all replicates as the best-fitting model. To further define the best-fitting parameters for this model, we ran another 100 inference replicates and determined the best-fit parameter combination based on the highest log-likelihood score.

As the demographic histories inferred by fastsimcoal2 and *δaδi* were fairly distinct (see Results below), we simulated data using fastsimcoal2 under the best-performing fastsimcoal2 models and the best-performing *δaδi* models to directly compare to the empirical data in order to further study model fit, keeping all simulated polymorphic sites and producing a single simulated SFS per replicate (’*-n*1 *-m -s*0 *--jobs --header --noarloutput -u -k* 6200000’). We simulated 100 replicates of the entire set of putatively neutral sites across the chromosomes (specifically, 22 autosomes were simulated with the same recombination and mutation rates used during inference and chromosome sizes of 81,997,209; 57,596,708; 59,482,194; 51,452,482; 56,421,937; 37,428,171; 57,377,548; 42,072,168; 32,858,669; 16,254,630; 13,862,685; 78,475,508; 36,722,319; 41,001,960; 32,326,613; 37,251,058; 16,847,824; 37,884,098; 19,023,431; 37,470,834; 27,914,525; and 13,844,647 bps). We then averaged the SFS from each replicate to produce the mean SFS for the best-performing fastsimcoal2 models and the best-performing *δaδi* models and visually compared the mean SFS for each model to the empirical SFS.

## Results

### Population genomic data for demographic inference

To infer the demographic dynamics of the coppery titi monkey, we deep-sequenced the genomes of six unrelated individuals housed at the CNPRC (Supplementary Table 2). After mapping the sequencing reads to the species-specific reference genome (PleCup_hybrid; Pfeifer et al. 2024), we called both biallelic variant and monoallelic invariant sites, limiting the dataset to putatively neutral, autosomal regions to circumvent the biasing effects of purifying and background selection on demographic inference (Ewing and Jensen 2014, 2016; Johri et al. 2020, 2021; Charlesworth and Jensen 2021, 2024). In total, 6 million variants were discovered in the accessible genome (Supplementary Table 3). Figure 1 provides a visual summary of this genome-wide, neutral data, showing Watterson’s θ and Tajima’s *D* for each autosome.

**Figure 1:**
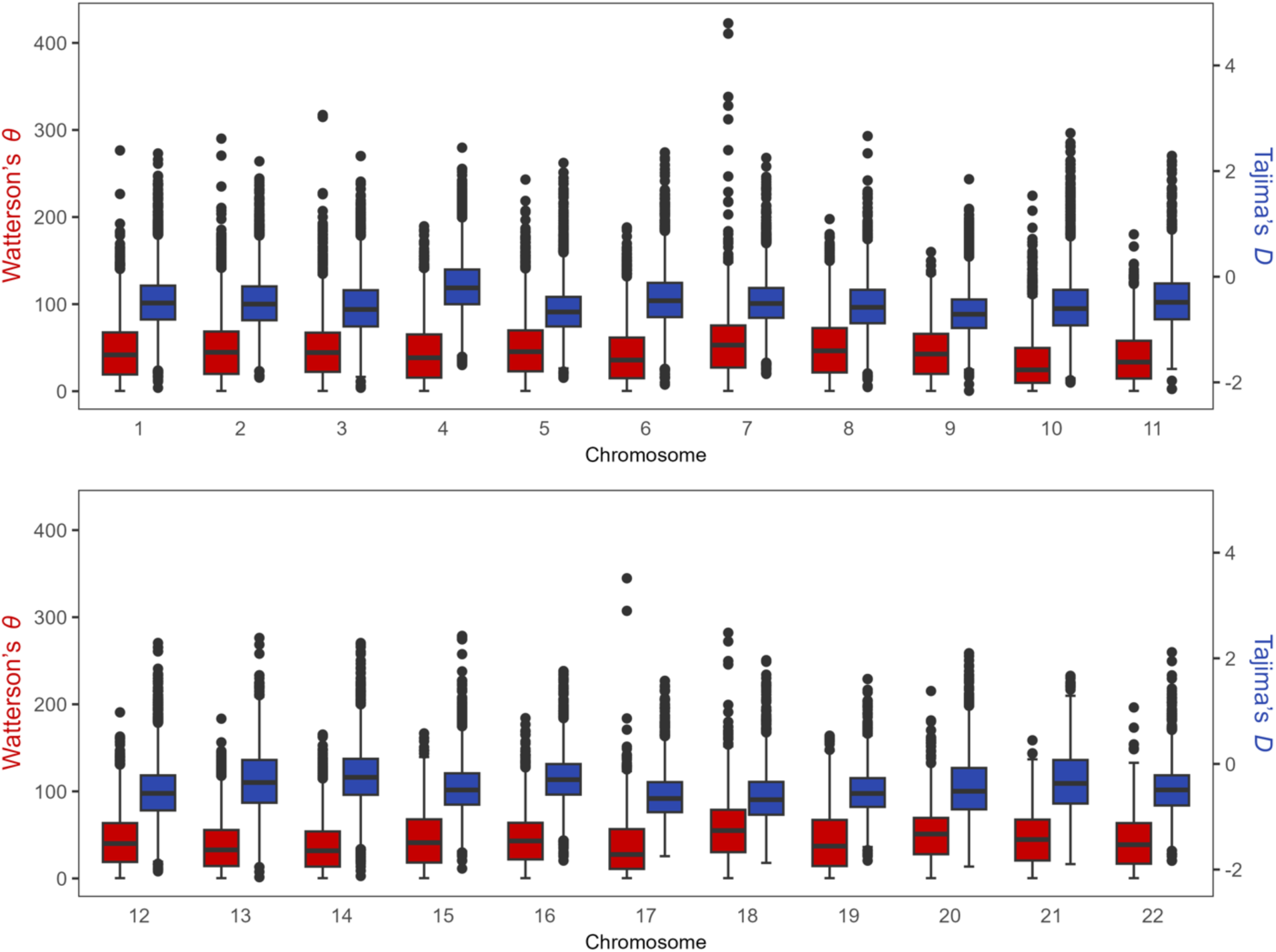
Summary of the genome-wide, neutral data being analyzed. The left axis provides the scaling for Watterson’s θ (plotted in red), the right axis provides the scaling for Tajima’s *D* (plotted in blue) – both based on 50 kb genomic windows – and the x-axis provides the chromosome being plotted.

### Estimating population structure

The fastSTRUCTURE analyses strongly concluded that all samples were collected from a single deme. The assignment of individuals to a specific deme was nearly complete with *Q* > 0.99999 ancestry components for all analyses. For both the full autosomal genome and individual autosomes, a value of *k* = 1 provided the greatest marginal likelihood (for the full autosomal genome, *L* = -1.102912 for *k* = 1; likelihood values for individual autosomes can be found in Supplementary Table 4); moreover, the full autosomal genome as well as most autosomes assigned all individuals to a single deme even for values of *k* greater than 1. Therefore, based on these analyses, our sampled individuals are likely of a single deme origin.

### Inferring the population size-change history

Based on this single deme result, we evaluated a variety of historical size change models (0 to 5 population size change events) using fastsimcoal2 (Excoffier et al. 2013), and the best parameterization was chosen based on the maximum likelihood scores of each replicate. The best performing models involved between 2 and 5 size change events (0-event: *L* = -19,827,291.914; 1-event: *L* = -19,755,162.832; 2-event: *L* = -19,747,605.136; 3-event: *L* = -19,747,259.099; 4-event: *L* = -19,747,248.682; 5-event: *L* = -19,747,239.622). Diagrams of the best-performing parameters for each of these models can be found in Figure 2 for the 3-event model, and in Supplementary Figure 1 for all other models (numerical parameter values are provided in Supplementary Table 5). Considering an alternative mutational scaling for the best-fitting model (1.07e-8/site/generation [supported by indirect estimation; Soni, Versoza, et al. 2026] instead of 4.88e-9/site/generation as used above [supported by direct estimation; Versoza et al. 2026a]), we observed a scaling down of estimated population sizes and more recent size change events (Supplementary Figure 2). This result is to be expected, given that changing the neutral mutation rate does not alter the SFS shape and thus does not alter the fit of the generalized model, but rather serves to re-scale the underlying parameters (e.g., population sizes scale smaller to keep the product of population size and mutation rate constant, and events scale more recently given the population size-based generation time scaling). Thus, it is important to be mindful of the impact of this underlying mutation rate uncertainty on the resulting demographic estimation; nonetheless, as direct pedigree-based mutation rate estimation is considered to be the gold-standard, it is fortunate to have such estimation available in this species.

**Figure 2:**
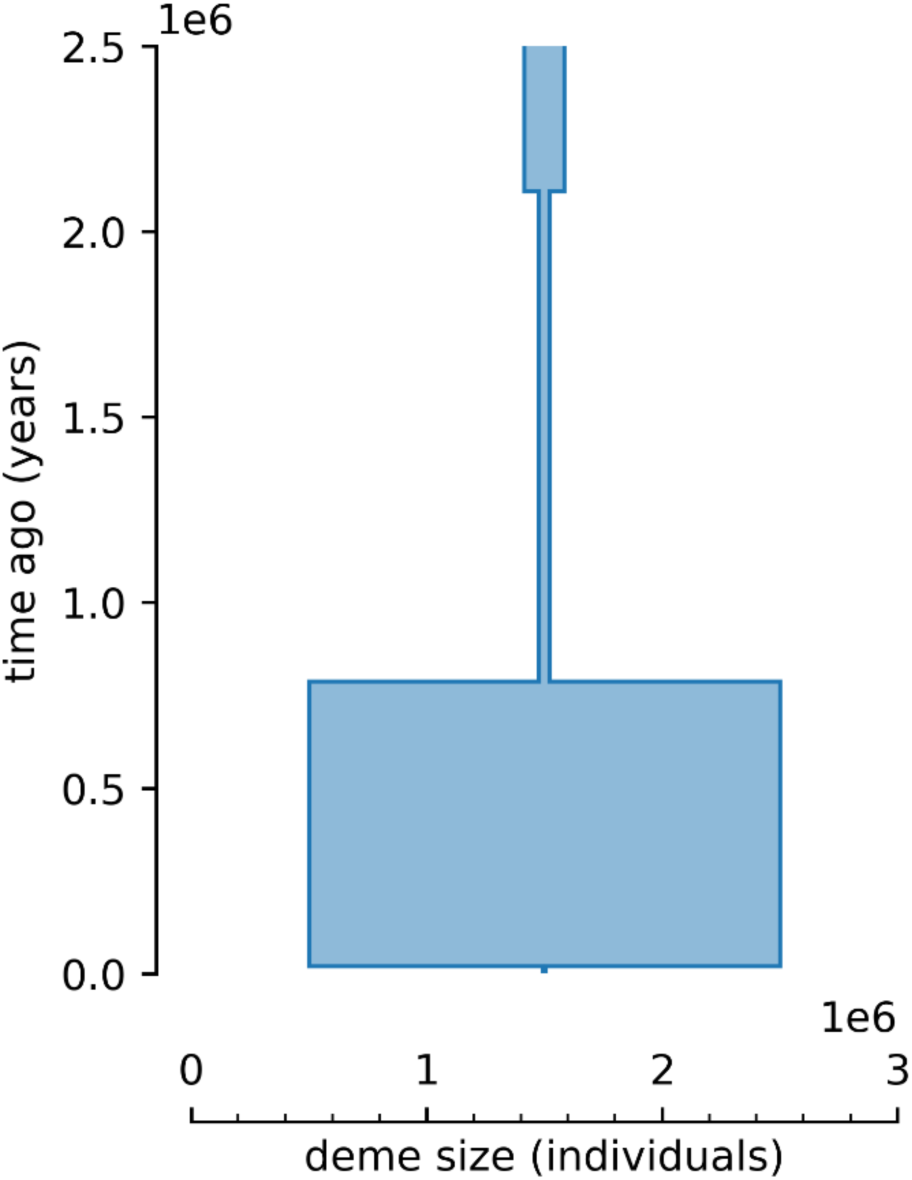
Diagram of best-performing fastsimcoal2 model, with 3 size-change events. The size of the population (in diploid individuals, scaled by 1e6) is represented by the width of the rectangles in the diagram, and the duration during which the population maintained that size (in years, scaled by 1e6) is indicated by the height of the rectangles in the diagram. Briefly, an ancestral population size of ∼170,000 individuals contracted to ∼46,000 individuals around 2.1 mya, grew to a population size of nearly 2,000,000 individuals within the past 1 mya, and contracted to ∼12,500 individuals within the past 20,000 years.

For comparative purposes, we also performed demographic inference using *δaδi* (Gutenkunst et al. 2009), an SFS-based neutral demographic estimator that infers population history via diffusion approximation. We initially tested seven single-population demographic models (see Material and Methods for more details), in order to identify the best-fitting model. The 5-epoch model (in which four instantaneous size changes occur) had the highest log-likelihood (-4,537.61), and the fewer parameters of the 3-epoch model (consisting of two instantaneous size changes) also provided a strong fit despite a lower log-likelihood (-19,500.89; and see Discussion below). Example diagrams of models are presented in Supplementary Figure 3 (specific numerical values scaled based on a mutation rate of 4.88e-9 /site/generation are available in Supplementary Table 6).

### Distinguishing between estimated demographic histories

As the composite likelihood scores emitted by fastsimcoal2 do not include any penalties for the use of additional parameters (each additional size change event requires two additional parameters), the relative improvement over simpler models, as well as the absolute difference from the observed composite likelihood, are important considerations when selecting the preferred model. The 3-event model is the simplest model within 50 units of the observed composite likelihood, with the additions of the 4-event and 5-event models resulting in likelihood scores with very minor improvements (Supplementary Table 7). Additionally, one would anticipate better performing models to consistently produce replicates that represent improvements over simpler models. The 1-event model produced 84.4% of replicates that outperform the constant size model, the 2-event model produced 43.0% of replicates that outperform the 1-event model, the 3-event model produced 5.8% of replicates that outperform the 2-event model, the 4-event model produced 0.6% of replicates that outperform the 3-event model, and the 5-event model produced 0.4% of replicates that outperform the 4-event model. As such, the 3-event model is the most complex model still resulting in >5% performance improvement amongst replicates. Finally, we compared the spread of the estimated parameters for better performing replicates using z-scores for the 0-, 1-, 2-, and 3-event models (Supplementary Figure 4). While the 0- and 1-event models are decidedly the most consistent in the resulting estimated parameters, the parameter estimates of the 3-event model exhibit comparatively less spread than the 2-event model. Taken together, these results thus suggest that the 3-event history is the best-performing fastsimcoal2 model, though if additional parameters are penalized more substantially, the 2-size change event model performs well with two fewer parameters. Notably, the general history of the 3-event and 2-event models are highly similar, suggesting strong growth over the past 1 million years followed by a very recent decline.

To compare the parameterized models of fastsimcoal2 and *δaδi*, we simulated SFS under each of the estimated demographic models and visually compared the simulated SFS to the empirically observed SFS. The mean simulated SFS of fastsimcoal2’s best performing models (2- event to 5-event models) all reasonably recapitulated the observed SFS; in contrast, the 3-epoch and 4-epoch models from *δaδi* resulted in SFS distinct from the observed SFS (Supplementary Figure 5). Thus, the 3-event model of fastsimcoal2 continues to appear most consistent with the observed data (Figure 3).

**Figure 3:**
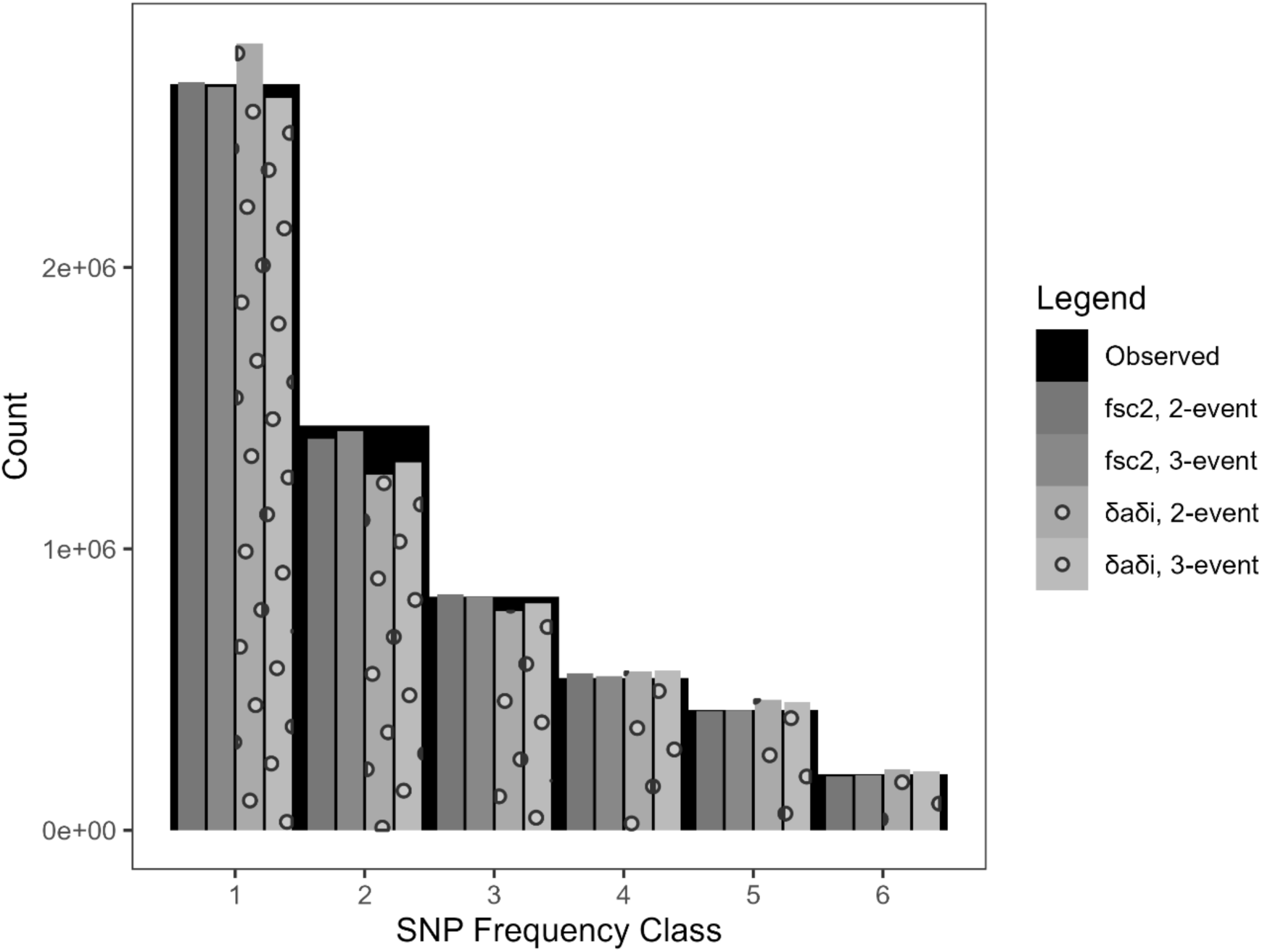
Mean simulated SFS for the two best fastsimcoal2 models (2-event and 3-event shown in solid gray bars, foreground) and two best *δaδi* models (shown in dotted gray bars, foreground) compared to the empirically observed SFS (black bar, background). SFS were simulated using fastsimcoal2 under the best parameters for each model. Diagrams of these models can be found in Figure 2 (fsc2, 3-event), Supplementary Figure 1 (fsc2, 2-event), and Supplementary Figure 3 (δaδi, 2-event and δaδi, 3-event). Mean simulated SFS for the 0-, 1-, and 4-event models parameterized by fastsimcoal2 can be found in Supplementary Figure 5.

## Discussion

Our results indicate that the sequenced individuals were sampled from a single population. While previous results have suggested population structuring (Byrne et al. 2016), it is most likely that all of our samples simply derived from a single deme and gene flow between / amongst demes is restricted, at least in the genetic ancestry of this sample. Alternatively, if there is sufficient gene flow amongst the demes, there may be no detectable genetic structuring. With regards to the ancestral history of population size change, in order to distinguish between our best performing models as derived from two commonly used inference approaches, we relied upon direct comparisons between predicted and empirically observed SFS. Scaling the timings of events from units of generations to years under these models (assuming a generation time of six years), a number of notable observations arise. Firstly, the bottleneck estimated at 2.1 mya by fastsimcoal2 corresponds with the previously rapid speciation occurring in the *moloch* group about 1.44 mya to 3.44 mya (Byrne et al. 2016, 2018; Byrne 2017). Moreover, the subsequent size changes correspond to major climatic shifts during the Pleistocene (Figure 4). Notably, the timing of a large population size expansion to nearly 2 million individuals is dated to about 786 kya, corresponding to the end of the Mid-Pleistocene Transition in which the glaciation cycles shifted from a 41,000- year periodicity to around a 100,000-year period; these longer cycles produced larger and more stable glacial sheets which would have led to overall dryer and cooler conditions (Clapperton 1992; Cook and Vizy 2006). Finally, the most recent estimated size change event is a relatively severe contraction dating to about 19 kya, which corresponds with the Pleistocene-Holocene boundary and the end of the Last Glacial Maximum (Hughes et al. 2013; Palacios et al. 2020).

**Figure 4:**
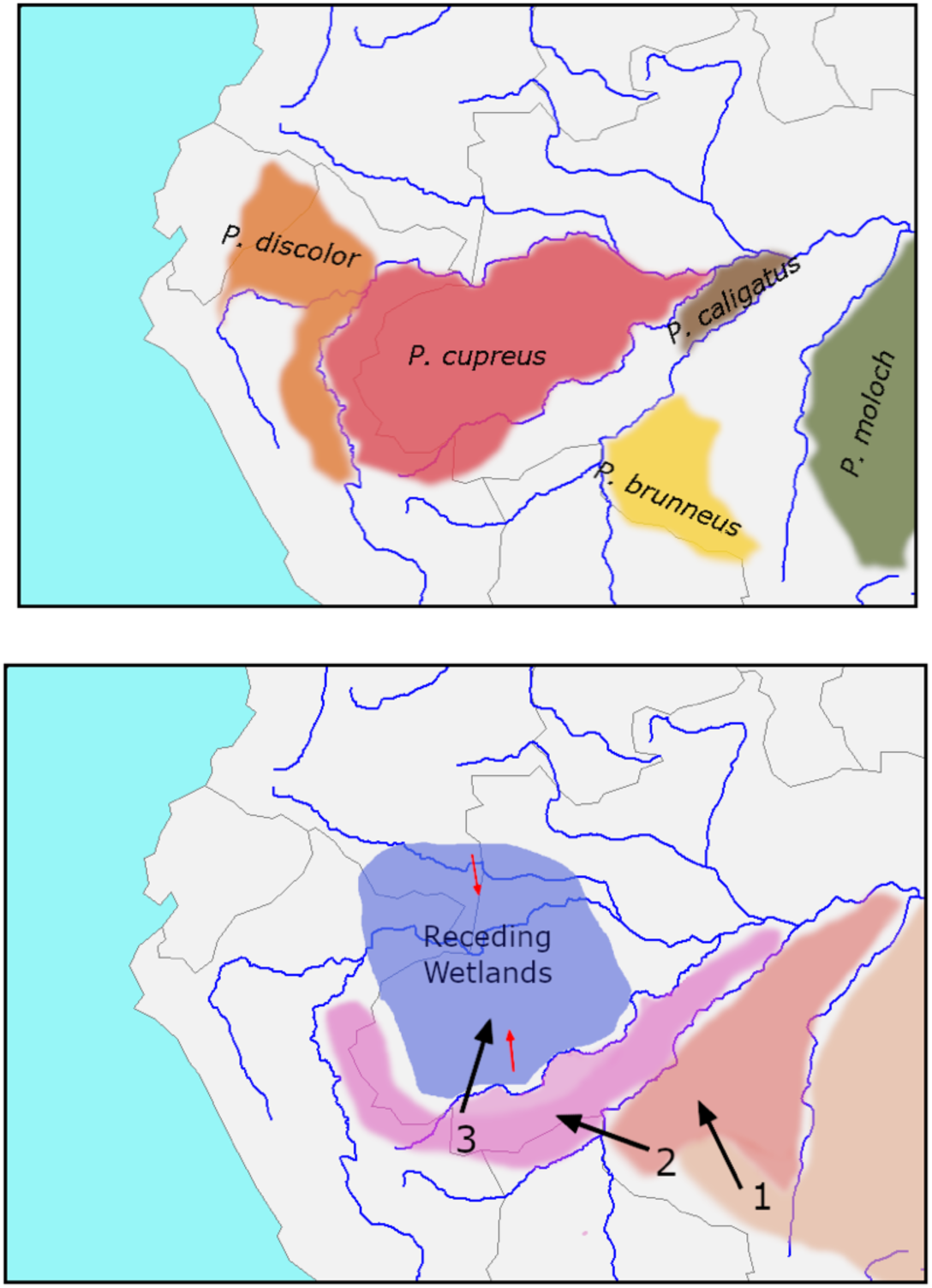
The biogeography underlying the modern and historical *P. cupreus* ranges. **Top panel**: Approximate current ranges of *P. moloch* and four species in the Western *moloch* group (*P. brunneus*, *P. caligatus*, *P. cupreus*, and *P. discolor*) based on data from the IUCN Red List (2025). Note that this is not an exhaustive depiction of all extant *Plecturocebus* species. **Bottom panel**: Diagram of the biogeographic model of speciation proposed by Byrne et al. (2018) for the Western *moloch* group. During the Pliocene, the final remnants of the mega-wetlands remaining from the Miocene Pebas and Acre systems (Hoorn et al. 2010) were receding and transitioning to *terra firme* rainforests (indicated in the diagram by the small red arrows in the “receding wetlands”), a process nearly finished by the early Pleistocene ∼2.5 mya. Accompanying this was migration into the new rainforest ecosystems by the common ancestors of the *moloch* group, (1) first expanding North and West leading to the split in the clade containing *P. miltoni* and *P. hoffmannsi* ∼2.24 mya. Following this, further expansion (2) led to the divergence between the Eastern and Western *moloch* groups ∼1.8-2.0 mya. Finally, as the ancestors of the Western *moloch* group reached the newest expansions of rainforest into the former mega-wetlands, (3) divergence between the members of the Western *moloch* group began ∼1.8 mya. Note that the rivers depicted are their current locations and not necessarily related to the ancient waterways that were present throughout this timespan.

Thus, these inferred population size changes appear to fit with the general pattern in the larger taxonomic group of range expansions and speciation following dryings of wetland ecosystems such as the Pebas system (Byrne 2017; Byrne et al. 2018), as well as with the current ecology of the coppery titi monkey relying on dry and/or seasonally wet terrestrial ecosystems. As global climate change leads to warmer weather and changes in precipitation patterns including increased river flooding (Almazroui et al. 2021; Alifu et al. 2022), this could lead to ecosystem changes detrimental to the coppery titi monkey, a species otherwise thought to be under little threat (Heymann et al. 2021). As such, an improved understanding of how past changes in climate relate to the population sizes of species like the coppery titi monkey sheds light on how the impacts of future climate change may be expected to affect species across the world, including those not currently considered vulnerable. Finally, aside from gaining novel insights into the population history of the species, this demographic null model will be useful for future genomic studies which may seek to quantify selective dynamics (as discussed in Johri et al. 2022), particularly given that this species is important as a model organism for the study of the neurophysiology underlying social attachment and monogamous pair bonds.

## ACKNOWLEDGEMENTS

DNA extraction, library preparation, and Illumina sequencing were conducted at the DNA Technologies and Expression Analysis Core at the UC Davis Genome Center (supported by NIH Shared Instrumentation Grant 1S10OD010786-01) and Novogene (Sacramento, CA, USA). Computations were performed on the Sol supercomputer at Arizona State University (Jennewein et al. 2023).

## References

Alifu H, Hirabayashi Y, Imada Y, Shiogama H. 2022. Enhancement of river flooding due to global warming. Sci Rep. 12(1):2–7. doi:10.1038/s41598-022-25182-6.

Almazroui M, Ashfaq M, Islam MN, Rashid IU, Kamil S, Abid MA, O’Brien E, Ismail M, Reboita MS, Sörensson AA, et al. 2021. Assessment of CMIP6 performance and projected temperature and precipitation changes over South America. Earth Syst Environ. 5(2):155–183. doi:10.1007/s41748-021-00233-6.

Baldwin MK, Krubitzer L. 2018. Architectonic characteristics of the visual thalamus and superior colliculus in titi monkeys. J Comp Neurol. 526(11):1750–1776. doi:10.1002/cne.24445

Bales KL, Ardekani CS, Baxter A, Karaskiewicz CL, Kusek JX, Lau AR, Savidge LE, Sayler KR, Witczak LR. 2021. What is a pair bond? Horm Behav. 136:105062. doi:10.1016/j.yhbeh.2021.105062

Bales KL, Mason WA, Catana C, Cherry SR, Mendoza SP. 2007. Neural correlates of pair-bonding in a monogamous primate. Brain Res. 1184(1):245–253. doi:10.1016/j.brainres.2007.09.087.

Browning BL, Tian X, Zhou Y, Browning SR. 2021. Fast two-stage phasing of large-scale sequence data. Am J Hum Genet. 108(10):1880–1890. doi:10.1016/j.ajhg.2021.08.005.

Byrne H. 2017. Evolutionary history and taxonomy of the titi monkeys (Callicebinae). University of Salford.

Byrne H, Lynch Alfaro JW, Sampaio I, Farias I, Schneider H, Hrbek T, Boubli JP. 2018. Titi monkey biogeography: parallel Pleistocene spread by *Plecturocebus* and *Cheracebus* into a post-Pebas Western Amazon. Zool Scr. 47(5):499–517. doi:10.1111/zsc.12300.

Byrne H, Rylands AB, Carneiro JC, Alfaro JWL, Bertuol F, da Silva MNF, Messias M, Groves CP, Mittermeier RA, Farias I, et al. 2016. Phylogenetic relationships of the New World titi monkeys (Callicebus): first appraisal of taxonomy based on molecular evidence. Front Zool. 13(1):10. doi:10.1186/s12983-016-0142-4.

Charlesworth B, Jensen JD. 2021. Effects of selection at linked sites on patterns of genetic variability. Annu Rev Ecol Evol Syst. 52(1):177–197. doi:10.1146/annurev-ecolsys-010621-044528.

Charlesworth B, Jensen JD. 2024. Population genetics. In: Scheiner SM, editor. Encyclopedia of biodiversity. 3rd ed. Amsterdam: Elsevier; p. 467–483.

Chintalapati M, Moorjani P. 2020. Evolution of the mutation rate across primates. Curr Opin Genet Dev. 62:58–64. doi: 10.1016/j.gde.2020.05.028.

Clapperton CM. 1993. Nature of environmental changes in South America at the Last Glacial Maximum. Palaeogeogr Palaeoclimatol Palaeoecol. 101(3–4):189–208. doi:10.1016/0031-0182(93)90012-8.

Conley AJ, Berger T, del Razo RA, Cotterman RF, Sahagún E, Goetze LR, Jacob S, Weinstein TAR, Dufek ME, Mendoza SP, et al. 2022. The onset of puberty in colony-housed male and female titi monkeys (*Plecturocebus cupreus*): possible effects of oxytocin treatment during peri-adolescent development. Horm Behav. 142:105157. doi:10.1016/j.yhbeh.2022.105157.

Cook KH, Vizy EK. 2006. South American climate during the Last Glacial Maximum: delayed onset of the South American monsoon. J Geophys Res Atmos. 111(2):1–21. doi:10.1029/2005JD005980.

Dolotovskaya S, Roos C, Heymann EW. 2020. Genetic monogamy and mate choice in a pair-living primate. Sci Rep. 10(1):20328. doi:10.1038/s41598-020-77132-9.

Ewing GB, Jensen JD. 2014. Distinguishing neutral from deleterious mutations in growing populations. Front Genet. 5(8):684–687. doi:10.3389/fgene.2014.00007.

Ewing GB, Jensen JD. 2016. The consequences of not accounting for background selection in demographic inference. Mol Ecol. 25(1):135–141. doi:10.1111/mec.13390.

Excoffier L, Dupanloup I, Huerta-Sánchez E, Sousa VC, Foll M. 2013. Robust demographic inference from genomic and SNP data. PLoS Genet. 9(10):e1003905. doi:10.1371/journal.pgen.1003905.

Excoffier L, Marchi N, Marques DA, Matthey-Doret R, Gouy A, Sousa VC. 2021. fastsimcoal2: demographic inference under complex evolutionary scenarios. Bioinformatics. 37(24):4882–4885. doi:10.1093/bioinformatics/btab468.

Falush D, Stephens M, Pritchard JK. 2003. Inference of population structure using multilocus genotype data: linked loci and correlated allele frequencies. Genetics. 164(4):1567– 1587. doi:10.1093/genetics/164.4.1567.

Feldman R. 2017. The neurobiology of human attachments. Trends Cogn Sci. 21(2):80–99. doi:10.1016/j.tics.2016.11.007.

Fischer EK, Nowicki JP, O’Connell LA. 2019. Evolution of affiliation: patterns of convergence from genomes to behaviour. Philos Trans R Soc B Biol Sci. 374(1777):20180242. doi:10.1098/rstb.2018.0242.

Freeman SM, Walum H, Inoue K, Smith AL, Goodman MM, Bales KL, Young LJ. 2014. Neuroanatomical distribution of oxytocin and vasopressin 1a receptors in the socially monogamous coppery titi monkey (*Callicebus cupreus*). Neuroscience. 273:12–23. doi:10.1016/j.neuroscience.2014.04.055.

Gutenkunst RN, Hernandez RD, Williamson SH, Bustamante CD. 2009. Inferring the joint demographic history of multiple populations from multidimensional SNP frequency data. PLoS Genet. 5(10):e1000695. doi:10.1371/journal.pgen.1000695.

Heymann EW, Calouro AM, de la Torre S, Vermeer J. 2021. *Plecturocebus cupreus* (amended version of 2018 assessment). Red List Threat Species. doi:10.2305/IUCN.UK.2021-1.RLTS.T127530593A192453653.en

Hoorn C, Wesselingh FP, ter Steege H, Bermudez MA, Mora A, et al. 2010. Amazonia through time: Andean uplift, climate change, landscape evolution, and biodiversity. Science. 330(6006):927–931. doi:10.1126/science.1194585.

Hughes PD, Gibbard PL, Ehlers J. 2013. Timing of glaciation during the last glacial cycle: evaluating the concept of a global “Last Glacial Maximum” (LGM). Earth-Science Rev. 125:171–198. doi:10.1016/j.earscirev.2013.07.003.

IUCN. 2025. The IUCN red list of threatened species. Version 2025-2. Available from: https://www.iucnredlist.org

Jennewein DM, Lee J, Kurtz C, Dizon W, Shaeffer I, Chapman A, et al. 2023. The Sol Supercomputer at Arizona State University. In Practice and Experience in Advanced Research Computing 2023: Computing for the Common Good (PEARC ’23). Association for Computing Machinery, New York, NY, USA, 296–301.

Jensen JD. 2023. Population genetic concerns related to the interpretation of empirical outliers and the neglect of common evolutionary processes. Heredity (Edinb). 130(3):109–110. doi:10.1038/s41437-022-00575-5.

Johri P, Aquadro CF, Beaumont M, Charlesworth B, Excoffier L, Eyre-Walker A, Keightley PD, Lynch M, McVean G, Payseur BA, et al. 2022. Recommendations for improving statistical inference in population genomics. PLoS Biol. 20(5):e3001669. doi:10.1371/journal.pbio.3001669.

Johri P, Charlesworth B, Jensen JD. 2020. Toward an evolutionarily appropriate null model: jointly inferring demography and purifying selection. Genetics. 215(1):173–192. doi:10.1534/genetics.119.303002.

Johri P, Pfeifer SP, Jensen JD. 2023. Developing an evolutionary baseline model for humans: jointly inferring purifying selection with population history. Mol Biol Evol. 40(5):1–14. doi:10.1093/molbev/msad100.

Johri P, Riall K, Becher H, Excoffier L, Charlesworth B, Jensen JD. 2021. The impact of purifying and background selection on the inference of population history: problems and prospects. Mol Biol Evol. 38(7):2986–3003. doi:10.1093/molbev/msab050.

Kuderna LFK, Ulirsch JC, Rashid S, Ameen M, Sundaram L, Hickey G, Cox AJ, Gao H, Kumar A, Aguet F, et al. 2024. Identification of constrained sequence elements across 239 primate genomes. Nature. 625(7996):735–742. doi:10.1038/s41586-023-06798-8.

Lau AR, Baxter A, He S, Loyant L, Ortiz-Jimenez CA, Bauman MD, Bales KL, Freeman SM. 2024. Age, pair tenure and parenting, but not face identity, predict looking behaviour in a pair-bonded South American primate. Anim Behav. 217:53–63. doi:10.1016/j.anbehav.2024.08.015.

Li H. 2013. Aligning sequence reads, clone sequences and assembly contigs with BWA-MEM. arXiv [Preprint] doi:10.48550/arXiv.1303.3997.

Marchi N, Kapopoulou A, Excoffier L. 2024. Demogenomic inference from spatially and temporally heterogeneous samples. Mol Ecol Resour. 24(1):53–63. doi:10.1111/1755-0998.13877.

Martin M. 2011. Cutadapt removes adapter sequences from high-throughput sequencing reads. EMBnet.journal 17:10. doi:10.14806/ej.17.1.200.

Mason WA. 1966. Social organization of the South American monkey, Callicebus moloch: a preliminary report. Tulane Stud Zool. 13:23–28.

Palacios D, Stokes CR, Phillips FM, Clague JJ, Alcalá-Reygosa J, Andrés N, Angel I, Blard PH, Briner JP, Hall BL, et al. 2020. The deglaciation of the Americas during the Last Glacial Termination. Earth-Science Rev. 203:103113. doi:10.1016/j.earscirev.2020.103113.

Pfeifer SP. 2017. From next-generation resequencing reads to a high-quality variant data set. Heredity (Edinb). 118(2):111–124. doi:10.1038/hdy.2016.102.

Pfeifer SP, Baxter A, Savidge LE, Sedlazeck FJ, Bales KL. 2024. *De novo* genome assembly for the coppery titi monkey (*Plecturocebus cupreus*): an emerging nonhuman primate model for behavioral research. Gen Biol Evol. 16(5):evae108. doi:10.1093/gbe/evae108/7675974.

Poh YP, Domingues V, Hoekstra HE, Jensen JD. 2014. On the prospect of identifying adaptive loci in recently bottleneck populations. PLoS One 9(11):e110579. doi:10.1371/journal.pone.0110579.

Pritchard JK, Stephens M, Donnelly P. 2000. Inference of population structure using multilocus genotype data. Genetics. 155(2):945–959. doi:10.1093/genetics/155.2.945.

Quinlan AR, Hall IM. 2010. BEDTools: a flexible suite of utilities for comparing genomic features. Bioinformatics. 26(6):841–842. doi:10.1093/bioinformatics/btq033.

Raj A, Stephens M, Pritchard JK. 2014. fastSTRUCTURE: variational inference of population structure in large SNP data sets. Genetics. 197(2):573–589. doi:10.1534/genetics.114.164350.

Soni V, Jensen JD. 2024. Temporal challenges in detecting balancing selection from population genomic data. G3 (Bethesda). 14(6):jkae069. doi:10.1093/g3journal/jkae069.

Soni V, Jensen JD. 2025. Inferring the demographic and selective histories from population genomic data using a two-step approach in species with coding-sparse genomes: an application to human data. G3 (Bethesda). 15(4):jkaf019. doi:10.1101/2024.09.19.613979.

Soni V, Terbot J, Versoza C, Pfeifer SP, Jensen JD. 2025. A whole-genome scan for evidence of positive and balancing selection in aye-ayes (*Daubentonia madagascariensis*) utilizing a well-fit evolutionary baseline model. G3 (Bethesda). 15(7):jkaf078. doi:10.1093/g3journal/jkaf078.

Soni V, Versoza CJ, Pfeifer SP, Jensen JD. 2025. Investigating the effects of chimerism on the inference of selection: quantifying genomic targets of purifying, positive, and balancing selection in common marmosets. Heredity 134:645–657.

Soni V, Versoza CJ, Terbot JW, Jensen JD, Pfeifer SP. 2025b. Inferring fine-scale mutation and recombination rate maps in aye-ayes (*Daubentonia madagascariensis*). Ecol Evol. 15:e72314.

Soni V, Versoza CJ, Spatola GJ, Bales KL, Jensen JD, Pfeifer SP. 2026. Inferring fine-scale rates of mutation and recombination in the coppery titi monkey (*Plecturocebus cupreus*). biorxiv, preprint.

Soni V, Versoza C, Vallender EJ, Jensen JD, Pfeifer SP. 2025a. Accounting for chimerism in demographic inference: reconstructing the history of common marmosets (*Callithrix jacchus*) from high-quality, whole-genome, population-level data. Mol Biol Evol. 42(6):msaf119. doi:10.1093/molbev/msaf119.

Terbot J, Calahorra-Oliart A, Versoza C, Shah D, Soni V, Pfeifer SP, Jensen JD. 2025b. Re-evaluating the demographic history of, and inferring the fine-scale recombination landscape for, wild Chinese rhesus macaques (*Macaca mulatta*). Am J Primatology. 87:e70088.

Terbot J, Soni V, Versoza C, Pfeifer SP, Jensen JD. 2025a. Inferring the demographic history of aye-ayes (*Daubentonia madagascariensis*) from high-quality, whole-genome, population-level data. Gen Biol Evol. 17(1):evae281. doi:10.1101/2024.11.08.622659.

Tran LAP, Pfeifer SP. 2018. Germline mutation rates in Old World monkeys. eLS, John Wiley & Sons, Ltd: Chichester. doi:10.1002/9780470015902.a0028242.

Van der Auwera G, O’Connor B. 2020. Genomics in the cloud: using Docker, GATK, and WDL in terra. Sebastopol (CA): O’Reilly Media.

Versoza CJ, Bales KL, Jensen JD, Pfeifer SP. 2026a. Characterizing rates and patterns of de novo germline mutations in coppery titi monkeys (*Plecturocebus cupreus*). biorxiv, preprint.

Versoza CJ, Bales KL, Jensen JD, Pfeifer SP. 2026b. Sex-specific landscapes of crossover and non-crossover recombination in coppery titi monkeys (*Plecturocebus cupreus*). biorxiv, preprint.

Versoza CJ, Lloret-Villas A, Jensen JD, Pfeifer SP. 2025. A pedigree-based map of crossovers and non-crossovers in aye-aye (*Daubentonia madagascariensis*). Gen Biol Evol. 17(5):evaf072. doi:10.1093/gbe/evaf072.

Versoza CJ, Weiss S, Johal R, La Rosa B, Jensen JD, Pfeifer SP. 2024. Novel insights into the landscape of crossover and non-crossover events in rhesus macaques (*Macaca mulatta)*. Gen Biol Evol. 16(1):evad223. doi:10.1093/gbe/evad223.

Zablocki-Thomas P, Rebout N, Karaskiewicz CL, Bales KL. 2023. Survival rates and mortality risks of *Plecturocebus cupreus* at the California National Primate Research Center. Am J Primatology. 85(10):e23531. doi:10.1002/ajp.23531.

