## Supplementary Materials for "Inferring the demographic history of coppery titi monkeys (*Plecturocebus cupreus*) from high-quality, whole-genome, population-level data"

**SUPPLEMENTARY MATERIAL**

| model | ***nu*** | | ***T*** | | ***nuB*** | | ***nuF*** | | ***TB*** | | ***TF*** | |
|---|---|---|---|---|---|---|---|---|---|---|---|---|
| | **lower** | **upper** | **lower** | **upper** | **lower** | **upper** | **lower** | **upper** | **lower** | **upper** | **lower** | **upper** |
| snm | - | - | - | - | - | - | - | - | - | - | - | - |
| 2-epoch | 0.01 | 10 | 0.001 | 5 | - | - | - | - | - | - | - | - |
| 3-epoch | - | - | - | - | 0.001 | 0.5 | 0.001 | 10 | 0.0001 | 0.05 | 0.0001 | 0.05 |
| growth | 0.001 | 10 | 0.001 | 5 | - | - | - | - | - | - | - | - |
| bottlegrowth | - | - | 0.001 | 5 | 0.001 | 0.5 | 0.001 | 10 | - | - | - | - |

| model | $t_1$ | | $t_2$ | | $t_3$ | | $t_4$ | | $N_1$ | | $N_2$ | | $N_3$ | | $N_4$ | |
|---|---|---|---|---|---|---|---|---|---|---|---|---|---|---|---|---|
| | **lower** | **upper** | **lower** | **upper** | **lower** | **upper** | **lower** | **upper** | **lower** | **upper** | **lower** | **upper** | **lower** | **upper** | **lower** | **upper** |
| 4-epoch | 0.01 | 10 | 0.01 | 10 | 0.01 | 10 | - | - | 0.001 | 5 | 0.001 | 5 | 0.001 | 5 | - | - |
| 5-epoch | 0.01 | 10 | 0.01 | 10 | 0.01 | 10 | 0.01 | 10 | 0.001 | 5 | 0.001 | 5 | 0.001 | 5 | 0.001 | 5 |

**Supplementary Table 1.** Lower and upper parameter bounds for each demographic model tested with *δaδi*: the standard neutral model (snm), as well as 2-epoch (a single instantaneous population size change), 3-epoch (two instantaneous population size changes), 4-epoch (three instantaneous population size changes), 5-epoch (four instantaneous size changes), growth (exponential population size change), and bottlegrowth (an instantaneous population size change followed by an exponential population size change) models. *nu* is the ratio of ancient to contemporary population size. *T* is the time in the past at which the size change occurred (in units of $2N_a$ generations, where $N_a$ is the ancestral population size). *nuB* is the ratio of the population size after instantaneous change to the ancient population size. *nuF* is the ratio of contemporary to the ancient population size. *TB* is the length of the bottleneck (in units of $2N_a$ generations). *TF* is the time since the bottleneck recovery (in units of $2N_a$ generations). For the 4- and 5-epoch models, the *t* values provide the timing of the size change event, and the *N* values the relative size change. If a parameter is not utilized in a given model, this is indicated by a “-“.

| NCBI ID | coverage |
|---|---|
| PleCup_01 | 43.0 |
| PleCup_02 | 39.4 |
| PleCup_03 | 38.8 |
| PleCup_04 | 56.6 |
| PleCup_05 | 42.9 |
| PleCup_06 | 37.9 |

**Supplementary Table 2.** Sample information including sequencing coverage (properly-paired autosomal reads only).

| | | putatively neutral | |
|---|---|---|---|
| **chromosome** | **length** | **# variant sites** | **# invariant sites** |
| 1 | 235,103,535 | 567,121 | 81,430,088 |
| 2 | 158,738,102 | 397,592 | 57,199,116 |
| 3 | 165,937,890 | 416,218 | 59,065,976 |
| 4 | 151,623,469 | 342,319 | 51,110,163 |
| 5 | 150,993,727 | 403,613 | 56,018,324 |
| 6 | 123,740,199 | 260,000 | 37,168,171 |
| 7 | 135,512,294 | 403,025 | 56,974,523 |
| 8 | 116,834,941 | 308,339 | 41,763,829 |
| 9 | 101,234,152 | 235,075 | 32,623,594 |
| 10 | 75,878,286 | 127,158 | 16,127,472 |
| 11 | 49,373,254 | 96,426 | 13,766,259 |
| 12 | 231,176,762 | 521,111 | 77,954,397 |
| 13 | 118,029,634 | 226,559 | 36,495,760 |
| 14 | 130,776,705 | 240,351 | 40,761,609 |
| 15 | 95,533,141 | 220,113 | 32,106,500 |
| 16 | 94,172,006 | 233,231 | 37,017,827 |
| 17 | 73,180,778 | 123,344 | 16,724,480 |
| 18 | 92,880,398 | 289,853 | 37,594,245 |
| 19 | 65,768,754 | 142,992 | 18,880,439 |
| 20 | 91,752,637 | 259,783 | 37,211,051 |
| 21 | 73,432,007 | 181,734 | 27,732,791 |
| 22 | 44,717,611 | 95,256 | 13,749,391 |
| $\sum$ or ∅ | **2,576,390,282** | **6,091,213** | **879,476,005** |

**Supplementary Table 3.** Summary of population genomic data used for demographic inference.

| chromosome | *k* | marginal likelihood |
|---|---|---|
| 1 | 1 | -1.102781154 |
| 1 | 2 | -1.102794203 |
| 1 | 3 | -1.102799147 |
| 1 | 4 | -1.102801801 |
| 1 | 5 | -1.102803481 |
| 2 | 1 | -1.099492001 |
| 2 | 2 | -1.099510169 |
| 2 | 3 | -1.099517072 |
| 2 | 4 | -1.099520783 |
| 2 | 5 | -1.099523134 |
| 3 | 1 | -1.085588794 |
| 3 | 2 | -1.085606206 |
| 3 | 3 | -1.085612817 |
| 3 | 4 | -1.085616371 |
| 3 | 5 | -1.085618622 |
| 4 | 1 | -1.137252612 |
| 4 | 2 | -1.169887528 |
| 4 | 3 | -1.137281437 |
| 4 | 4 | -1.137285713 |
| 4 | 5 | -1.137288422 |
| 5 | 1 | -1.083092217 |
| 5 | 2 | -1.083110135 |
| 5 | 3 | -1.083116939 |
| 5 | 4 | -1.083120598 |
| 5 | 5 | -1.083122916 |
| 6 | 1 | -1.109385799 |
| 6 | 2 | -1.109412764 |
| 6 | 3 | -1.109423047 |
| 6 | 4 | -1.109428587 |
| 6 | 5 | -1.109432100 |
| 7 | 1 | -1.108846879 |
| 7 | 2 | -1.108864817 |
| 7 | 3 | -1.108871633 |
| 7 | 4 | -1.108875298 |
| 7 | 5 | -1.108877619 |
| 8 | 1 | -1.095654068 |
| 8 | 2 | -1.095677084 |
| 8 | 3 | -1.095685846 |
| 8 | 4 | -1.095690563 |
| 8 | 5 | -1.095693552 |
| 9 | 1 | -1.074410665 |
| 9 | 2 | -1.124492695 |
| 9 | 3 | -1.074451579 |
| 9 | 4 | -1.074457669 |
| 9 | 5 | -1.074461533 |
| 10 | 1 | -1.093497183 |
| 10 | 2 | -1.149639708 |
| 10 | 3 | -1.093569597 |
| 10 | 4 | -1.144533221 |
| 10 | 5 | -1.150501391 |

| | | |
|---|---|---|
| 11 | 1 | -1.100772536 |
| 11 | 2 | -1.139158542 |
| 11 | 3 | -1.139185420 |
| 11 | 4 | -1.100880193 |
| 11 | 5 | -1.100889151 |
| 12 | 1 | -1.095241499 |
| 12 | 2 | -1.095255620 |
| 12 | 3 | -1.095260973 |
| 12 | 4 | -1.095263848 |
| 12 | 5 | -1.095265668 |
| 13 | 1 | -1.125956726 |
| 13 | 2 | -1.125987363 |
| 13 | 3 | -1.125999065 |
| 13 | 4 | -1.126005373 |
| 13 | 5 | -1.126009375 |
| 14 | 1 | -1.132384222 |
| 14 | 2 | -1.132413222 |
| 14 | 3 | -1.132424295 |
| 14 | 4 | -1.132430262 |
| 14 | 5 | -1.132434046 |
| 15 | 1 | -1.105810580 |
| 15 | 2 | -1.105842052 |
| 15 | 3 | -1.105854073 |
| 15 | 4 | -1.105860554 |
| 15 | 5 | -1.105864666 |
| 16 | 1 | -1.127010759 |
| 16 | 2 | -1.141839448 |
| 16 | 3 | -1.127051969 |
| 16 | 4 | -1.141857351 |
| 16 | 5 | -1.141861249 |
| 17 | 1 | -1.084425396 |
| 17 | 2 | -1.138525803 |
| 17 | 3 | -1.084499885 |
| 17 | 4 | -1.084511057 |
| 17 | 5 | -1.084518160 |
| 18 | 1 | -1.083172478 |
| 18 | 2 | -1.083196855 |
| 18 | 3 | -1.083206141 |
| 18 | 4 | -1.083211141 |
| 18 | 5 | -1.083214310 |
| 19 | 1 | -1.097024759 |
| 19 | 2 | -1.148785295 |
| 19 | 3 | -1.151662885 |
| 19 | 4 | -1.097099424 |
| 19 | 5 | -1.145346869 |
| 20 | 1 | -1.105843432 |
| 20 | 2 | -1.143790435 |
| 20 | 3 | -1.105880709 |
| 20 | 4 | -1.105886253 |
| 20 | 5 | -1.105889769 |
| 21 | 1 | -1.127882203 |
| 21 | 2 | -1.128205407 |
| 21 | 3 | -1.127934175 |

| | | |
|---|---|---|
| 21 | 4 | -1.128227967 |
| 21 | 5 | -1.127946867 |
| 22 | 1 | -1.104309552 |
| 22 | 2 | -1.154615298 |
| 22 | 3 | -1.104404192 |
| 22 | 4 | -1.157483783 |
| 22 | 5 | -1.104427497 |
| all autosomes | 1 | -1.102912093 |
| all autosomes | 2 | -1.102913503 |
| all autosomes | 3 | -1.102914028 |
| all autosomes | 4 | -1.102914307 |
| all autosomes | 5 | -1.102914483 |

**Supplementary Table 4.** fastSTRUCTURE results for the full autosomal genome and individual autosomes for values of *k* (number of demes) from 1 to 5.

| *model* | $N_{ANC}$ | $T_1$ | $N_1$ | $T_2$ | $N_2$ | $T_3$ | $N_3$ | $T_4$ | $N_4$ | $T_5$ | $N_5$ |
|---|---|---|---|---|---|---|---|---|---|---|---|
| *fsc2-0* | 115,697 | – | – | – | – | – | – | – | – | – | – |
| *fsc2-1* | 1,650 | 1,860,516 | 181,719 | – | – | – | – | – | – | – | – |
| *fsc2-2* | 46,437 | 971,940 | 2,709,188 | 185,388 | 81,529 | – | – | – | – | – | – |
| ***fsc2-3*** | **169,079** | **2,107,008** | **45,656** | **786,090** | **1,999,796** | **18,960** | **12,338** | **–** | **–** | **–** | **–** |
| *fsc2-4* | 172,443 | 2,042,346 | 43,330 | 823,938 | 924,184 | 428,196 | 663,758 | 3,864 | 3,238 | – | – |
| *fsc2-5* | 337,368 | 2,440,314 | 206,222 | 2,031,000 | 502,541 | 1,015,830 | 3,344 | 933,696 | 879,124 | 18,282 | 13,092 |

**Supplementary Table 5.** The estimated best parameter values for each of the models tested with fastsimcoal2 (0 to 5 population size change events). Time is given in units of years before present and population sizes are given in units of diploid individuals. The best model presented in this study (3-size change; fsc2-3) is shown in bold. The time points (*T*) signify the timing of the corresponding population sizes change events from the previous population size *N* to the subsequent population size *N*. These parameters correspond to Figure 2 for the 3-size change model, and Supplementary Figure 1 for all other models. If a parameter is not utilized in a given model, this is indicated by a "-".

| model | $N_{ANC}$ | $T_1$ | $N_1$ | $T_2$ | $N_2$ | $T_3$ | $N_3$ | $T_4$ | $N_4$ |
|---|---|---|---|---|---|---|---|---|---|
| *δaδi-1* | 25,832 | 1,543,871 | 182,554 | – | – | – | – | – | – |
| *δaδi-2* | 2,505,581 | 3,598,625 | 2,872 | 1,837,423 | 182,632 | – | – | – | – |
| *δaδi-3* | 1,617,146 | 53,042,551 | 17,451 | 1,569,150 | 217,624 | 69,705 | 92,528 | – | – |
| *δaδi-4* | 1,928,775 | 18,003,131 | 147,808 | 1,669,540 | 24,602 | 1,248,573 | 272,393 | 158,430 | 118,498 |
| *δaδi-growth* | 40,564 | 2,430,703 | 217,846 | – | – | – | – | – | – |
| *δaδi-bottlegrowth* | 411,141 | 4,586,649 | 9,730 | – | 217,348 | – | – | – | – |

**Supplementary Table 6.** The estimated best parameter values for each of the models tested with *δaδi*: the standard neutral model (*δaδi*-1), as well as 2-epoch (a single instantaneous population size change), 3-epoch (two instantaneous population size changes), 4-epoch (three instantaneous population size changes), growth (exponential population size change), and bottlegrowth (an instantaneous population size change followed by an exponential population size change) models. Time is given in units of years before present and population sizes are given in units of diploid individuals. The time points (*T*) signify the timing of the corresponding population sizes change events from the previous population size *N* to the subsequent population size *N*. The parameters of *δaδi*-2 and *δaδi*-3 correspond to the diagrams in Supplementary Figure 3. If a parameter is not utilized in a given model, this is indicated by a "-".

| *model* | *k* | maxL | ΔL | % improvement |
|---|---|---|---|---|
| *fsc2-0* | 1 | -19827291.91 | 80052.29 | 100.0 |
| *fsc2-1* | 3 | -19755162.83 | 7923.21 | 84.4 |
| *fsc2-2* | 5 | -19747605.14 | 365.51 | 43.0 |
| ***fsc2-3*** | **7** | **-19747259.10** | **19.48** | **5.8** |
| *fsc2-4* | 9 | -19747248.68 | 9.06 | 0.6 |
| *fsc2-5* | 11 | -19747239.62 | 0.00 | 0.4 |
| *observed* | N/A | -19747222.53 | (-17.09) | N/A |

**Supplementary Table 7.** Comparison of likelihood scores, and the fraction of replicates with improvements relative to the next simplest model, for models analyzed with fastsimcoal2 (0 to 5 population size change events). The best model presented in this study (3-size change; fsc2-3) is shown in bold.

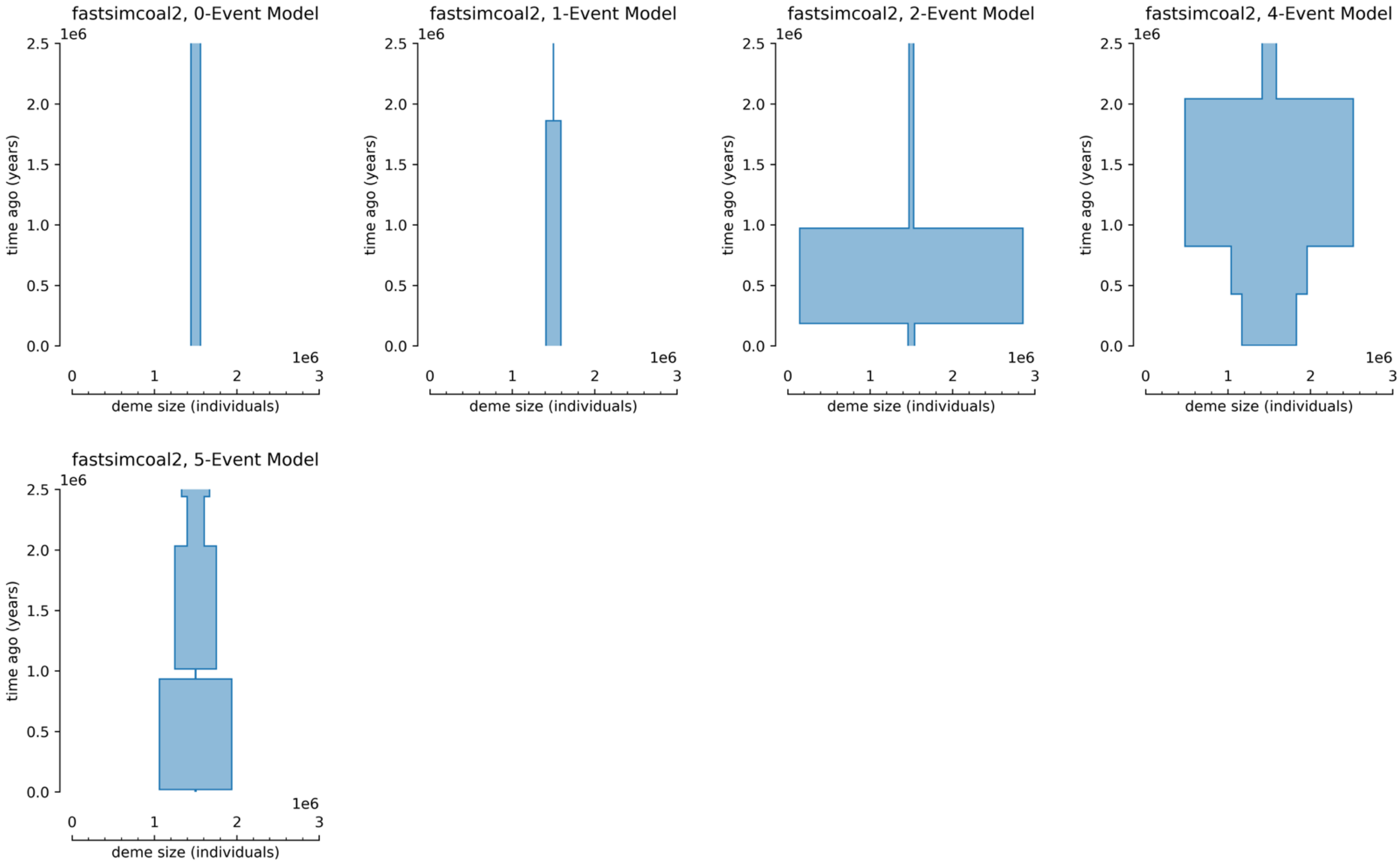


**Supplementary Figure 1.** Diagrams of alternative fastsimcoal2 models (0, 1, 2, 4, and 5 population size change events). Population size (in diploid individuals, scaled by 1e6) is represented by the width of the rectangles in each diagram, and the duration of that size (in years, scaled by 1e6) is represented by the height of the rectangles in the diagrams.

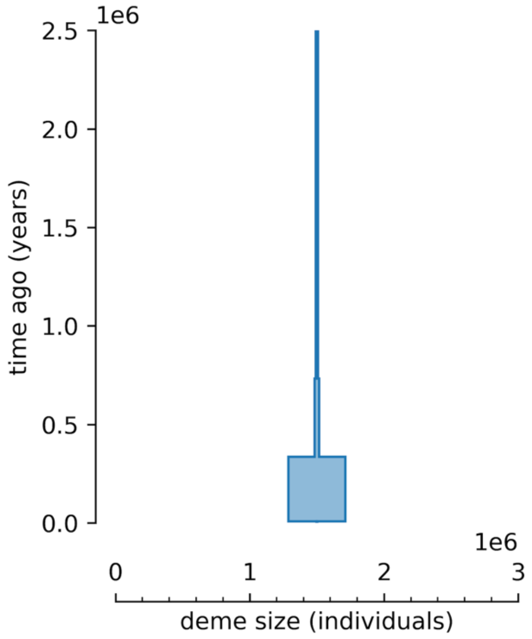


**Supplementary Figure 2.** Diagram of the impact of a higher underlying assumption of mutation rate on the best-fitting model presented in Figure 2. Population size (in diploid individuals, scaled by 1e6) is represented by the width of the rectangles in each diagram, and the duration of that size (in years, scaled by 1e6) is represented by the height of the rectangles in the diagrams.

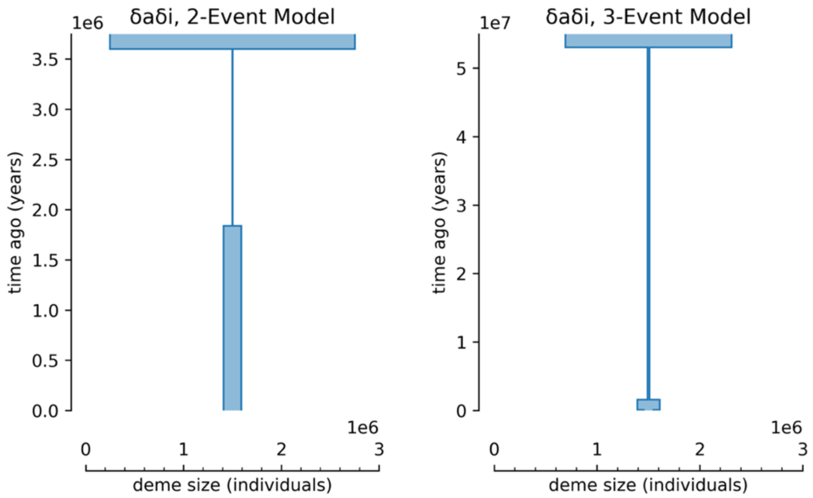


**Supplementary Figure 3.** Diagrams of 2-event and 3-event *δaδi* models. Population size (in diploid individuals, scaled by 1e6) is represented by the width of the rectangles in each diagram, and the duration of that size (in years, scaled by 1e6 (left) and 1e7 (right)) is represented by the height of the rectangles in the diagrams.

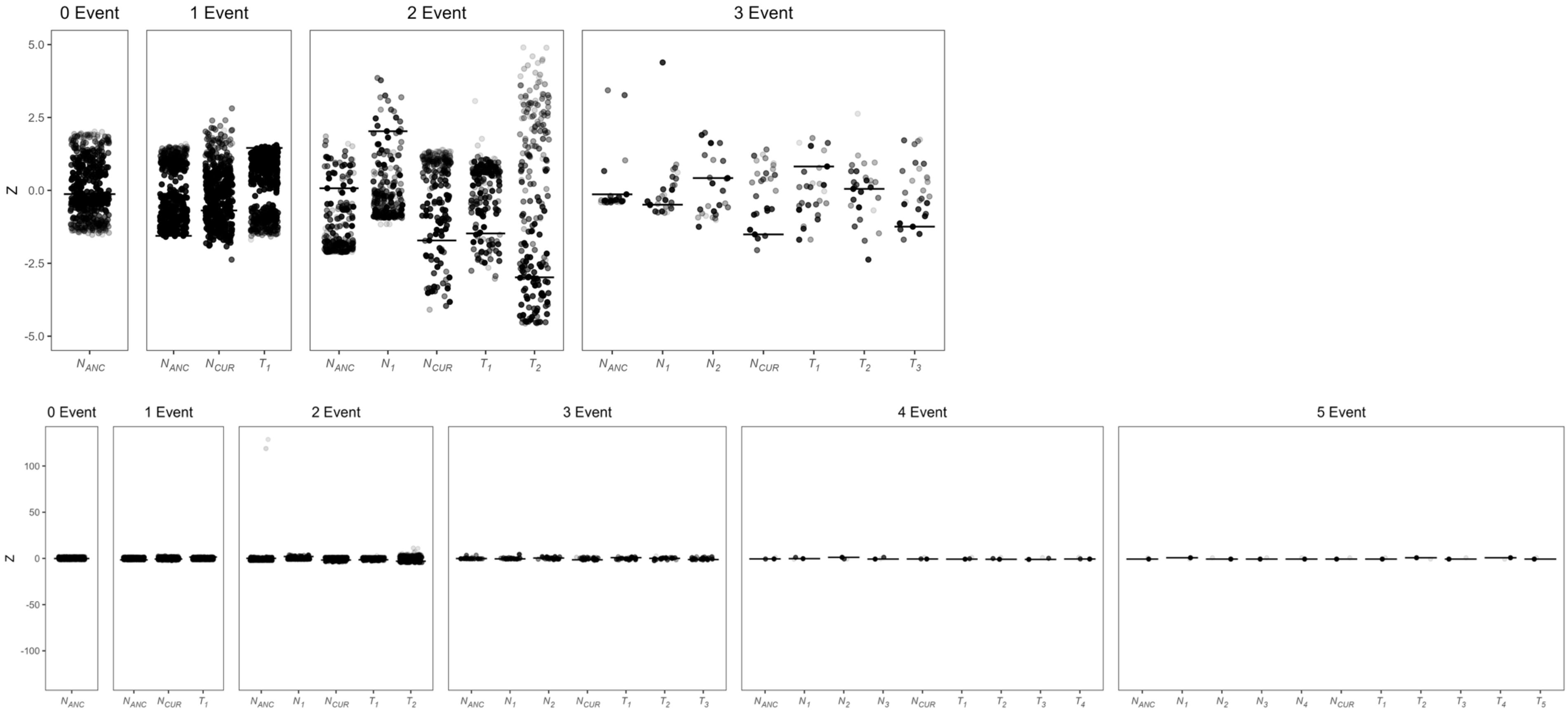


**Supplementary Figure 4.** z-scores for estimated parameter values for each replicate of the models analyzed with fastsimcoal2 (top panel), which represented improvements (based on likelihood scores) over the next simplest model. Note that this figure excludes the 4- and 5-event models, and is zoomed in such that outliers are not visible (the zoomed-out version is provided in the bottom panel). The opacity of points corresponds to the likelihood scores within each model (i.e., the darkest point for each parameter corresponds to the best-fitting parameters of each model). The horizontal bars for each model indicate the z-score for the best-fitting parameters.

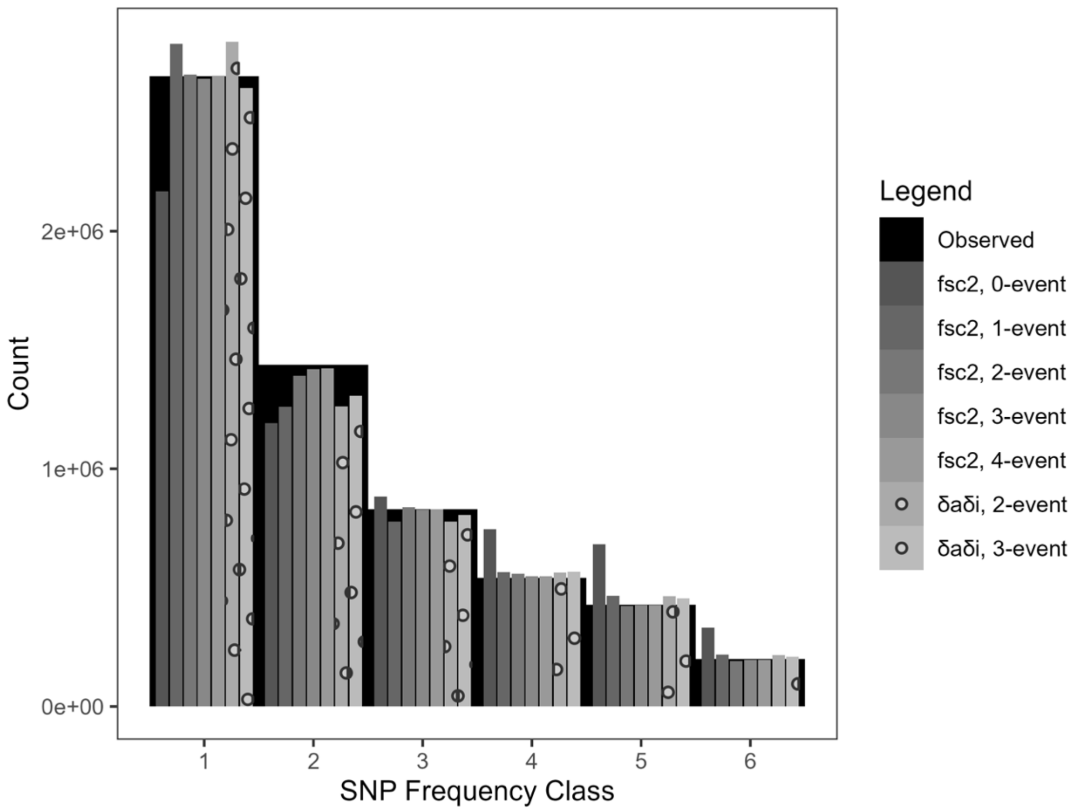


**Supplementary Figure 5.** Simulated SFS for the five fastsimcoal2 models (0 to 5 population size change events shown in solid gray, foreground) and two *δaδi* models (2-event and 3-event shown in dotted gray, foreground), compared to the empirically observed SFS (black bar, background). SFS were simulated using fastsimcoal2 under the best parameters for each model. Details of the parameters for these models can be found in Supplementary Tables 5 and 6 (and see Figure 2 as well as Supplementary Figures 1 and 3 for diagrams of these models).
